# No evidence for somatosensory attenuation during action observation of self-touch

**DOI:** 10.1101/2021.02.08.430193

**Authors:** Konstantina Kilteni, Patrick Engeler, Ida Boberg, Linnea Maurex, H. Henrik Ehrsson

## Abstract

The discovery of mirror neurons in the macaque brain in the 1990s triggered investigations on putative human mirror neurons and their potential functionality. The leading proposed function has been action understanding: accordingly, we understand the actions of others by ‘simulating’ them in our own motor system through a direct matching of the visual information to our own motor programs. Furthermore, it has been proposed that this simulation involves the prediction of the sensory consequences of the observed action, similar to the prediction of the sensory consequences of our executed actions. Here, we tested this proposal by quantifying somatosensory attenuation behaviorally during action observation. Somatosensory attenuation manifests during voluntary action and refers to the perception of self-generated touches as less intense than identical externally generated touches because the self-generated touches are predicted from the motor command. Therefore, we reasoned that if an observer simulates the observed action and, thus, he/she predicts its somatosensory consequences, then he/she should attenuate tactile stimuli simultaneously delivered to his/her corresponding body part. In three separate experiments, we found a systematic attenuation of touches during executed self-touch actions, but we found no evidence for attenuation when such actions were observed. Failure to observe somatosensory attenuation during observation of self-touch is not compatible with the hypothesis that the putative human mirror neuron system automatically simulates the observed action. In contrast, our findings emphasize a sharp distinction between the motor representations of self and others.

## Introduction

Mirror neurons are a class of neurons observed in the macaque brain that discharge both when the monkey executes a goal-directed action, such as grasping, and when the monkey observes another agent performing the same or a similar action (di Pellegrino et al. 1992; Gallese et al. 1996; Rizzolatti and Craighero 2004; Rizzolatti and Sinigaglia 2010). These neurons are different from motor neurons that discharge only during goal-directed actions and from ‘canonical’ neurons that discharge also during the presentation of three-dimensional objects (Oztop et al. 2013; Giese and Rizzolatti 2015). The mirror neurons were initially discovered in area F5 in the premotor cortex (di Pellegrino et al. 1992; Gallese et al. 1996), but neurons with mirror properties have also been reported in other areas, including the primary motor cortex (Vigneswaran et al. 2013), the inferior parietal lobule (Fogassi et al. 2005; Bonini et al. 2010), the lateral intraparietal area (Shepherd et al. 2009), and the medial frontal cortex (Yoshida et al. 2011) (see also (Kilner and Lemon 2013).

The discovery of mirror neurons in the macaque brain immediately triggered the question of whether human mirror neurons exist. In contrast to macaque research, systematic electrophysiological recordings in humans are not frequently feasible; therefore, neuroimaging research focused on the detection of active brain regions — increases in the synaptic activity of large neuronal populations rather than increased discharge rates of single neurons — that are responsive to both action observation and action execution. Early neuroimaging experiments indeed revealed an overlap between the action execution and action observation networks, including the supplementary motor area, the dorsal premotor cortex, the supramarginal gyrus and the superior parietal lobule (Grezes and Decety 2001) (see also (Caspers et al. 2010)). A meta-analysis of studies on action observation and/or action execution that attributed their findings to the putative human mirror neuron system (Molenberghs et al. 2012) revealed a number of brain areas displaying ‘mirror properties’, including the inferior parietal lobule, inferior frontal gyrus and the ventral premotor cortex – areas that are considered homologous to the areas in the monkey cortex containing mirror neurons (Rizzolatti and Craighero 2004). In relation, a recent meta-analysis (Hardwick et al. 2018) revealed the coactivation of the supplementary motor area, the ventral and dorsal premotor cortex and the inferior parietal lobule during both action observation and action execution. However, neural activity during action observation can represent other non-mirroring processes, such as visual recognition, motion perception and working memory. In addition, the activation of areas during both action observation and action execution might be driven by visuomotor neurons other than mirror neurons (Dinstein et al. 2008; Oztop et al. 2013). In response to this criticism, experiments investigating neural adaptation to repeated stimuli presentation were conducted to test whether areas with putative mirror neurons show adaptation to repeated actions independently of whether these were observed or executed. However, these studies yielded mixed results (Dinstein et al. 2007; Chong et al. 2008; Lingnau et al. 2009). Finally, using single-cell recordings, Mukamel and colleagues (2010) recorded activity from neurons that discharged during both the observation and the execution of actions. These neurons were located at the supplementary motor area, the hippocampus and parahippocampal gyrus and the entorhinal cortex, while the classic mirror neuron locations could not be tested due to the limitations in the placement of the recording electrodes.

Mirror neurons have been assigned a wide variety of functions, ranging from action understanding and imitation to empathy, emotion recognition and language learning (for discussion, see (Rizzolatti and Craighero 2004; Heyes 2010a; Rizzolatti and Sinigaglia 2010; Cook et al. 2014)). With respect to action understanding, it has been postulated that when observing actions performed by others, the motor system of the observer ‘resonates’ with the visual experience of the action (di Pellegrino et al. 1992; Rizzolatti and Fadiga 1998; Rizzolatti et al. 2001). This motor resonance is accomplished through a *simulation* process, during which the observer simulates the seen action using his/her own motor system involving his/her mirror neurons (Gallese and Goldman 1998; Decety and Grèzes 2006; de Vignemont and Haggard 2008). According to the so-called ‘direct-matching hypothesis’(Iacoboni et al. 1999; Rizzolatti et al. 2001), the visual representation of the observed action is directly matched onto a motor representation in the observer’s brain in an automatic fashion (Rizzolatti and Sinigaglia 2010; Uithol et al. 2011). Mirror neurons represent this mapping between the sensory representation of the observed action and the related motor program. Through this automatic simulation, we understand the observed actions and their goal, as well as the intention of the observed agent ((Umiltà et al. 2001; Rizzolatti and Sinigaglia 2010), but see (Hickok 2009; Lingnau et al. 2009; Uithol et al. 2011)).

How mirror neurons perform this mapping of the visual experience to the motor program during action observation remains unknown. Computational models of human motor control have been used to provide a possible mechanism. Accordingly, the central nervous system uses inverse models to transform intended sensory states into motor commands and forward models to predict the sensory consequences of these motor commands (Wolpert et al. 1995, 1998, 2001; Miall and Wolpert 1996; Wolpert and Kawato 1998; Wolpert and Flanagan 2001; Davidson and Wolpert 2005). Moreover, a parallel architecture of multiple pairs of inverse and forward models can allow the system to switch between different pairs that best fit the current sensorimotor context (Haruno et al. 2001; Wolpert et al. 2003). Internal simulation loops of motor control have been proposed to serve mirror functions, such as inferring the goal of the observed action (Oztop et al. 2006, 2013; Hurley 2008; Imamizu 2010). Accordingly, during action observation, the matching of the visual representation of the observed action to the associated motor command would correspond to inverse modeling (Arbib and Rizzolatti 1996; Oztop et al. 2013). Iacoboni et al. (1999; 2005) proposed that mirror neurons are also involved in forward modeling; connections in the macaque brain from the superior temporal sulcus to the posterior parietal cortex and then to mirror neurons in area F5 could represent the inverse model, while connections from area F5 to the posterior parietal cortex and then to the superior temporal sulcus could represent the predicted visual consequences of the movements –, i.e., the forward model. In an extension of this proposal, Miall (2003) included the cerebellum and the posterior parietal cortex in both the inverse and the forward internal model circuitries. Demiris and Johnsson (2003) proposed a distributed simulation-based architecture in which the observer’s inverse models estimate the motor commands necessary to achieve the state of the observed agent. These motor commands are fed to forward models to predict the next sensory state of the observed person. The predicted sensory consequences are then compared with the received sensory consequences of the observed person, and errors will determine which pair of inverse and forward models better fits the observed action (Wolpert et al. 2003). Within this computational framework, mirror neuron activity can be seen as the predicted sensory consequences of the observed action, i.e., the outcome of the forward models (Oztop et al. 2006, 2013).

Here, we aimed to experimentally assess the recruitment of forward models during action observation by quantifying the phenomenon of somatosensory attenuation. Somatosensory attenuation refers to the perception that self-generated touches feel less intense than external touches of the same intensity (Blakemore et al. 1998, 1999; Shergill et al. 2003, 2014; Bays et al. 2005; Walsh et al. 2011; Kilteni and Ehrsson 2017a, b, 2020b, a; Kilteni et al. 2018, 2019, 2020) (see also (Lalouni et al. 2020) for recent findings in pain attenuation). Computational motor control theories suggest that attenuation occurs because self-generated tactile sensations can be predicted by the forward model using a copy of the motor command, i.e., an efference copy, in contrast to external stimuli (Blakemore et al. 2000; Wolpert and Flanagan 2001; Bays and Wolpert 2008). Therefore, we reasoned that if, during action observation, the forward models of the observer are recruited to simulate the observed action, then the observation of an actress moving her right index finger to touch her left finger should enable the observer to predict the touch on the actress’s left index finger. Because of this somatosensory prediction, a tactile stimulus simultaneously delivered to the observer’s left index finger would be attenuated. We performed three separate experiments to test this hypothesis. In all three experiments, we failed to detect any reliable evidence for somatosensory attenuation during action observation. Our data are, therefore, not compatible with the hypothesis that a key function of the putative human mirror neuron system is to simulate the observed action using the same sensorimotor computational mechanisms that are used for the control of movement.

## Materials and Methods

### Participants

After providing written informed consent, 30 participants (15 women and 15 men, 29 right-handed and 1 ambidextrous) aged 19–36 years participated in Experiment 1, 32 participants (16 women and 16 men, 30 right-handed and 2 ambidextrous) aged 19–33 years participated in Experiment 2 and 24 participants (12 women and 12 men, 24 right-handed) aged 18–36 years participated in Experiment 3.

The sample sizes of all studies were set to thirty (30) before data collection commenced based on our previous studies using the same methods (Kilteni et al. 2019, 2020; Kilteni and Ehrsson 2020a). To ensure a counterbalanced order of conditions in all experiments, the sample size was increased to 32 for Experiments 2 and 3. Unfortunately, however, due to the coronavirus disease 2019 (COVID19) restrictions in human testing, data collection for Experiment 3 had to be stopped after 24 participants.

In all experiments, handedness was assessed using the Edinburgh Handedness Inventory (Oldfield 1971). All experiments were approved by the Swedish Ethical Review Authority (no. 2019-03355).

### Experiment 1

#### Methods

In Experiment 1, participants performed a modified version of the force-matching task (Shergill et al. 2003). The participants sat at a table and rested their left hands palm up, with their index fingers on a molded support (**Figure 1**). In each trial, a constant force was applied on the pulp of their relaxed left index fingers (*applied* force) from a cylindrical probe (25 mm height) with a flat aluminum surface (20 mm diameter) attached to a lever controlled by a DC electric motor (Maxon EC Motor EC 90 flat; manufactured in Switzerland). A force sensor (FSG15N1A, Honeywell Inc., USA; diameter, 5 mm; minimum resolution, 0.01 N; response time, 1 ms; measurement range, 0–15 N) was placed inside the probe to measure the applied forces. Immediately after receiving the force, the participants were asked to generate a force that matched the intensity of the applied force (*matched force*). In all conditions, the participants moved the wiper of a 13 cm slide potentiometer with their right hands to match the forces (**Figure 1**). The slider controlled the force output on the participants’ left index fingers. The lower limit of the slider (left extreme) corresponded to 0 N and the maximum (right extreme) to 5 N. Each trial started with the slider at 0 N. Both the applied and the matched forces had a duration of 3 s. Each condition included 36 trials, with each applied force level (1, 1.5, 2, 2.5, 3 and 3.5 N) pseudorandomly presented six times.

**Figure 1.**
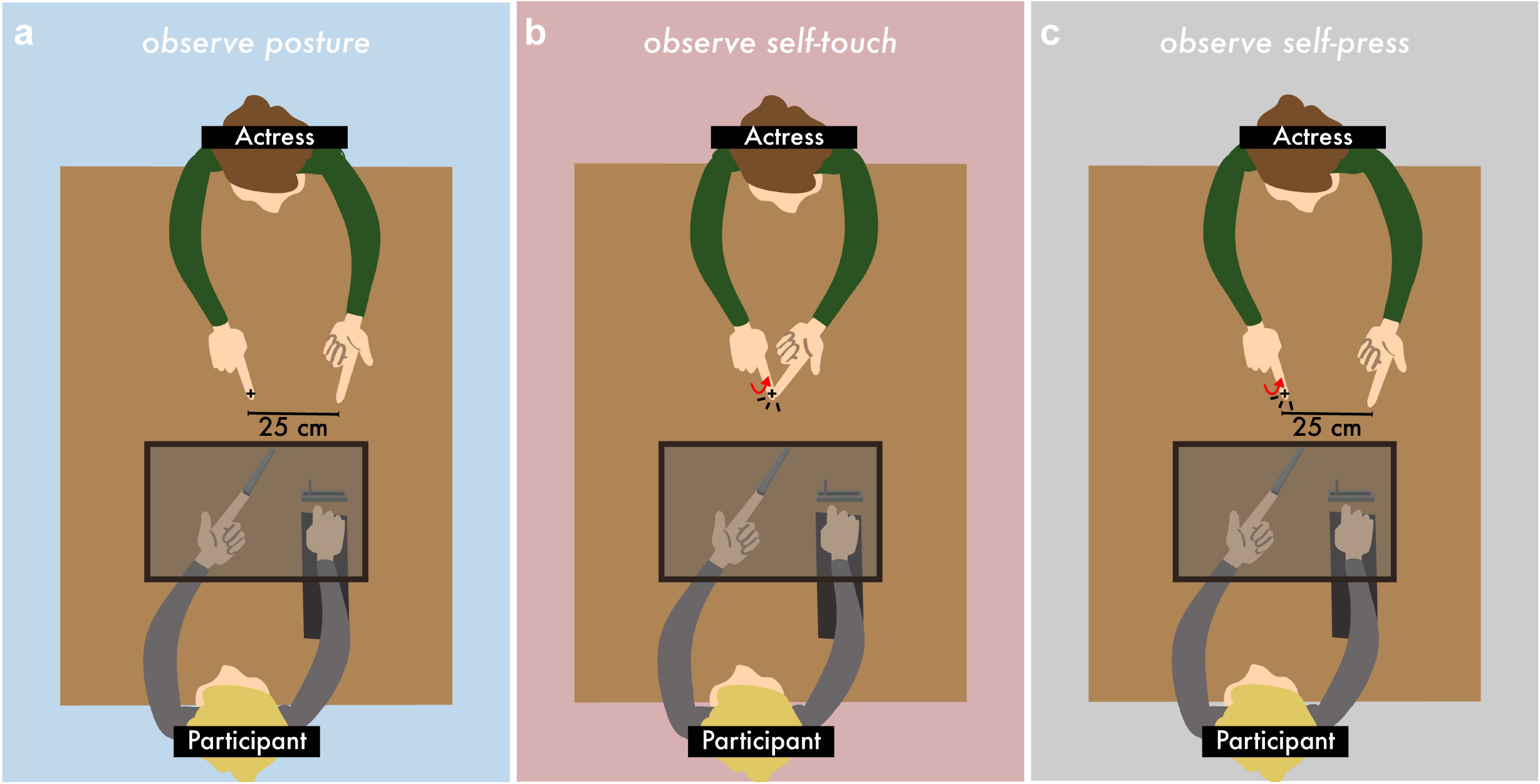
Experimental conditions in the force-matching task (Experiment 1). In all conditions, participants received a force on the pulp of their left index finger from a motor. **(a)** In the *observe posture* condition, the participants simultaneously observed a static posture of the actress’s hands. **(b)** In the *observe self-touch* condition, the participants simultaneously observed the actress performing a reaching movement to touch and press against her left index finger. The forces the actress applied to her finger matched the forces the participants were receiving at the same time. **(c)** In the *observe self-press* condition, the participants simultaneously observed the actress performing a reaching movement to touch and press against the table. As in (b), the forces the actress applied to the table matched the forces the participants were receiving at the same time. In all conditions, the participants reproduced the applied force by moving a slider with their right hand. The equipment and the participants’ hands were occluded with a screen. Although the screen is seen transparent for illustration purposes, it was actually opaque. Fixation points are denoted by a cross (+). Red arrows in (**b**) and (**c**) illustrate the starting and ending positions of the actress’ movement (movement extent ~3 cm).

Auditory signals indicated the onset and offset of the periods of the applied and matched forces, respectively. In all conditions, the equipment and the participants’ hands were occluded from view by a screen. In addition, participants wore headphones through which white noise was administered to preclude sounds produced by the motor to interfere with the task. The loudness of the white noise allowed participants to hear the auditory signals clearly. No feedback was provided to the participants concerning their performance in the force-matching task.

Three experimental conditions were presented in a counterbalanced order. All conditions involved the presence of an actress sitting opposite to the participant at the other side of a table (width x length, 70 × 120 cm) (**Figure 1**). Both the actress’s hands were placed on the table in full view of the participant. In all conditions, the actress’s left hand was placed palm up, and her right hand was placed palm down. Both her index fingers extended as if pointing, while the other fingers were flexed to curl under the palm.

In the *observe posture* condition, the actress placed her right index finger on a marked position on the table (which served as the fixation point) and her left index finger on another marked position at 25 cm to the left from her right index finger (**Figure 1a**). The actress did not move her hands during this condition and remained as motionless as possible throughout the trials. Upon the auditory cue, participants were asked to fixate on the actress’s right index finger (that remained motionless) while they simultaneously received the applied force from the motor. Following this, they reproduced the force with the slider as described above.

In the *observe self-touch* condition, the actress placed her left index finger on the marked position on the table (same as the fixation point) and her right index finger next to it (~3 cm to the right). Upon the auditory cue, the actress performed a fast reaching movement involving lifting her right upper arm, moving it leftwards and touching the pulp of her left index finger with the pulp of her right index finger (**Figure 1b**). Participants observed the actress’s right index finger reaching and pressing against her left one while they simultaneously received the applied force from the motor on the pulps of their left index fingers. Once the applied force finished, the actress placed her right index finger back to the table. Next, the participants reproduced the force with the slider.

A hidden force sensor placed underneath the actress’s left index finger measured her applied forces. The actress (but not the participants) received online visual feedback of her forces to match the applied forces participants were simultaneously receiving from the motor. That is, if participants received a force of 2 N on their left index finger, the actress aimed to press 2 N on her left index finger with her right index finger. We reasoned that by doing so, the observed action would optimally match the received sensory input in how forceful the action looked and thereby facilitate its central simulation and associated sensory attenuation.

In the *observe self-press* condition, the actress had her index fingers separated by 28 cm, similar to the *observe posture* condition, with her right index finger being next to the hidden sensor (~3 cm to the right). Upon the auditory cue, the actress performed a fast reaching movement with her right arm and hand to touch and press her right index finger against the table where the hidden sensor was placed (**Figure 1c**). As in the *observe self-touch* condition, the actress aimed to match the forces the participants were simultaneously receiving.

In all conditions, the fixation point of the participants was always at the same depth position (**Figure 1**). The initial arm postures of the participant and the actress were similar across conditions. In both observation conditions, the participants passively observed the actress without moving. The actress avoided eye contact with the participants except brief glances every 8-10 trials – once the participants had matched the force and before the next applied force started – to ensure that the participant fixated on the actress’s finger. The order of the conditions was fully counterbalanced.

After the end of all conditions, participants were asked whether they felt as if the actress was delivering the forces on their finger. We asked this question to control for any unspecific influences in somatosensory perception driven by who the participants perceived to deliver the force: the motor or the actress. All but one subject reported that the forces were controlled by the motor and not by the actress. Therefore, we did not ask this question in the subsequent two experiments.

#### Statistical analysis

The mean of the matched force data recorded from the participants’ left index finger sensors was calculated at 2500–3000 ms after the ‘go’ signal. These are the last 500 ms of the time participants had to match the applied forces (3000 ms); thus, they reflected their final estimation. Visual inspection ensured that the participants had not released the sensor during this interval. We then averaged the mean matched forces across the six repetitions of each force level. The same was done for the data recorded from the sensor of the actress’s left index finger. All data were extracted using Python (version 3.7) and analyzed using R (R version 3.4.4) (Core Team 2018) and JASP (JASP and JASP Team 2019). We used a repeated-measures analysis of variance (ANOVA) and paired t-tests since the data satisfied normality (Shapiro-Wilk test). In addition, a Bayesian factor analysis using default Cauchy priors with a scale of 0.707 was carried out for all parametric tests to provide information about the level of support for the null hypothesis compared to the alternative hypothesis (*BF*_*01*_) given the data.

#### Hypothesis

Previous experiments have shown that when participants use the slider to reproduce the forces, they accurately match the applied forces, i.e., they show no attenuation (Shergill et al. 2003; Wolpe et al. 2016; Kilteni and Ehrsson 2017a, b). Therefore, we expected to see an accurate performance during the *observe posture* condition, which served as a control condition. Importantly, we hypothesized that, during the observation of self-touch (*observe self-touch*), participants would simulate the reach-to-press movement of the actress in their own motor system and predict its somatosensory consequences (i.e., self-touch). This prediction should then attenuate the perceived intensity of the touch that is simultaneously delivered on the participants’ left index fingers. Therefore, when using the slider, participants would match it with a lower force. Finally, the *observe self-press* condition served as an additional control condition. This condition included the observation of a reach-to-press movement, but it did not have any somatosensory consequences for the left index finger, i.e., the actress presses against the table and not her left index finger. Therefore, we reasoned that participants would simulate the observed movement, but this simulation would not affect the perception of touch on their left index finger since there were no predicted tactile sensations in that finger. Finally, although we did not hypothesize any sex effects, we checked for their presence in a post hoc analysis given that the observed agent was always female.

### Experiment 2

#### Methods

As in Experiment 1, participants were asked to place their left index fingers inside the molded support with the palm up. The participants’ right forearms comfortably rested on top of a set of sponges next to their left hands. In each trial, the motor delivered two taps (the test tap and the comparison tap in **Figure 2a-d**) on the pulp of the participants’ left index fingers, and participants were asked to verbally indicate which tap was stronger. An auditory ‘go’ signal indicated the onset of the trial and the onset of the response period.

**Figure 2.**
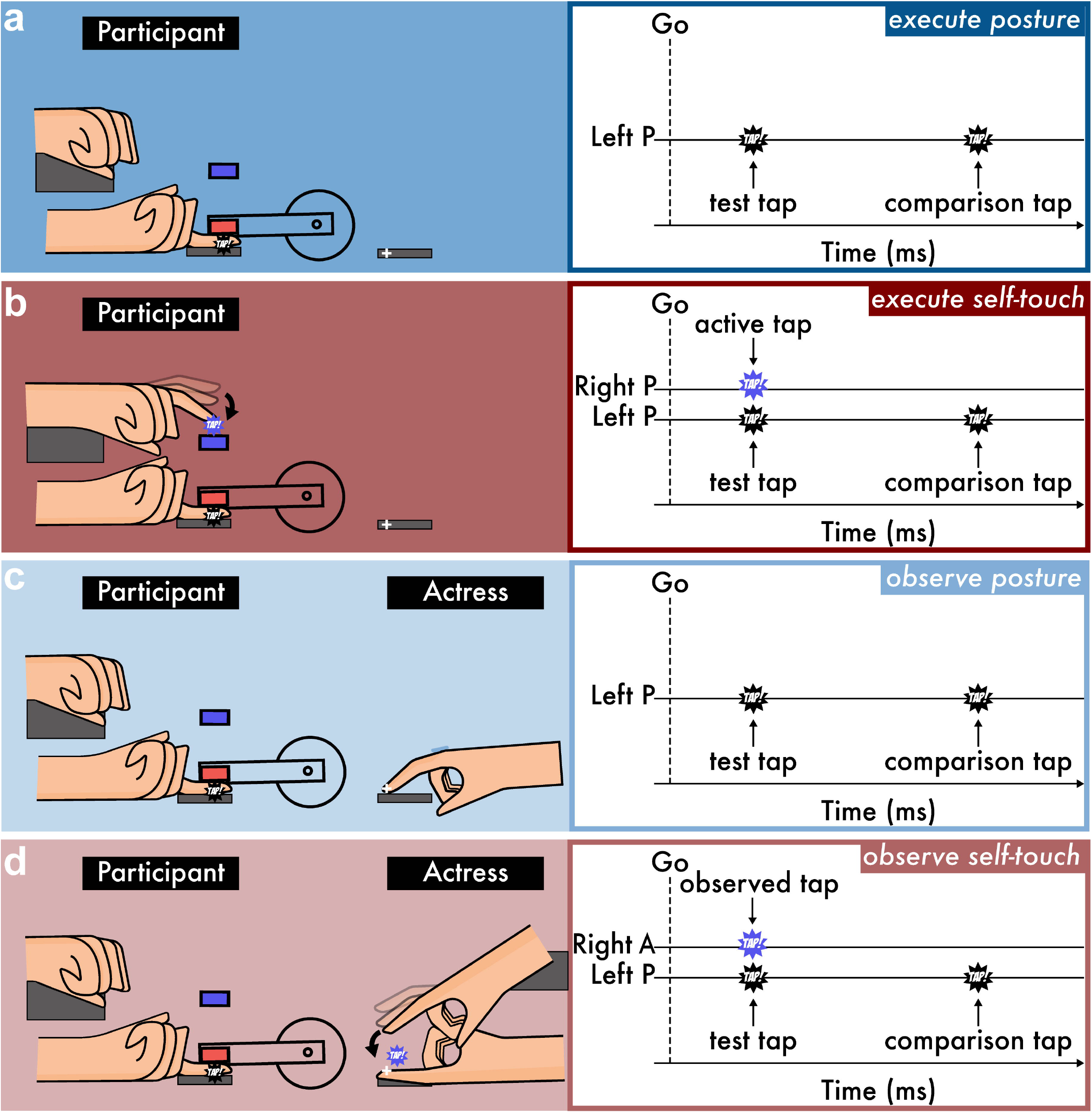
Experimental conditions in the force-discrimination task (Experiment 2). In all conditions, participants received two taps (test and comparison tap) on the pulp of their left index finger from the motor, and they had to verbally indicate which was stronger: the first or the second tap. **(a)** In the *execute posture* condition, the participants kept a relaxed posture while receiving the two taps. **(b)** In the *execute self-touch* condition, the participants triggered the test tap by actively tapping with their right index finger a force sensor (indicated in blue) placed just above their left index finger. **(c)** In the *observe posture* condition, the participants kept a relaxed posture but now fixated on the actress’s right index finger, which remained still while receiving the two taps. **(d)** In the *observe self-touch* condition, the actress triggered the test tap by actively tapping with her right index finger a hidden force sensor that was placed below her left index finger. Fixation points are denoted by a cross (+). Right P and left P represent the right and left index fingers of the participant, respectively, while right A represents the right index finger of the actress.

Four experimental conditions were grouped in two sessions (**Figure 2**): the ‘execution’ session, which did not involve the actress and consisted of two conditions (*execute posture* and *execute self-touch*), and the ‘observation’ session, which involved the actress and consisted of two conditions (*observe posture* and *observe self-touch*). The order of the sessions as well as the order of the conditions within each session were fully counterbalanced.

In the *execute posture* condition (**Figure 2a**), participants remained relaxed while the two taps were applied to their left index fingers after the auditory cue. This served as a control condition. In the *execute self-touch* condition (**Figure 2b**), participants triggered the test tap on their left index fingers by actively tapping with their right index fingers a force sensor placed just above their left index finger after the auditory cue. This active tap triggered the test tap with an intrinsic delay of ~35 ms. In both motor execution conditions, a mark placed on the table served as the fixation point of the participants (**Figure 2a-b**).

In the *observe posture* condition (**Figure 2c**), participants were asked to fixate on the actress’s right index finger, which remained still, while the two taps were applied to their left index fingers after the auditory cue. The left hand of the actress was placed 25 cm to the left of her right hand and remained still. This condition served as a control condition, controlling for the effect of simply observing an actor and her hands. Finally, in the *observe self-touch* condition (**Figure 2d**), the actress’s left index finger was placed on top of the mark on the table (palm up), and her right index finger (palm down) was positioned 3 cm to the right of her left index finger. Upon the auditory cue, the actress actively tapped the pulp of her left index finger with the pulp of her right index finger. A hidden force sensor placed just underneath her left index finger detected the active tap and triggered the test tap with an intrinsic delay of ~35 ms. In both observation conditions, the participants passively observed the actress without moving. The fixation point was the actress’s left index finger, which was placed on top of the mark on the table (**Figure 2c-d**). The initial arm postures of the participant and the actress were similar across conditions.

Each condition involved 70 trials. The test tap was set to 2 N, while the intensity of the comparison tap was systematically varied among seven different force levels (1, 1.5, 1.75, 2, 2.25, 2.5 or 3 N). The order of the taps was the same in all experimental conditions. The two taps had a 100-ms duration, and the delay between them was random (800 – 1500 ms). In the *execute posture* and *observe posture* conditions, the test tap was applied 500 ms after the auditory cue.

In the *execute self-touch* condition, participants were asked to tap, neither too weakly nor too strongly, with their right index finger, as if tapping the screen of their smartphone. Similar instructions were given to the actress in the *observe self-touch* condition.

As before, the actress avoided eye contact with the participants, except brief glances every 8-10 trials – once the participants had given response – to ensure that they fixated on her finger. During the ‘execution’ session, where the actress was not seen, participants were verbally reminded to fixate on the mark placed on the table every 8-10 trials. Again, in all conditions, the equipment and the participants’ hands were occluded from view by a screen, participants wore headphones and no feedback was ever provided to them concerning their performance.

#### Statistical analysis

In each condition, we fitted the participants’ responses with a generalized linear model using a logistic function (**Equation 1**), similar to previous studies (Bays et al. 2005, 2006; Kilteni et al. 2019, 2020; Kilteni and Ehrsson 2020b). Two parameters of interest were extracted: the point of subjective equality (PSE), which represents the intensity at which the test tap felt as strong as the comparison tap (p = 0.5) and which quantifies the attenuation, and the just noticeable difference (JND), which reflects the participants’ sensitivity in force discrimination.

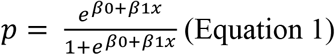

Thirteen force trials (13 of 8960, 0.14%) were rejected because either the intensity of the test tap (2 N) was not applied correctly or the responses were not recorded. Before fitting the responses, the values of the applied comparison taps were binned to the closest value with respect to their theoretical values (1, 1.5, 1.75, 2, 2.25, 2.5 or 3 N). To contrast the conditions of interest, we performed paired t-tests or Wilcoxon signed-rank tests depending on whether the data satisfied normality (Shapiro-Wilk test). In addition, a Bayesian factor analysis using default Cauchy priors with a scale of 0.707 was carried out for all parametric tests to provide information about the level of support for the null hypothesis compared to the alternative hypothesis (*BF*_*01*_) given the data.

#### Hypothesis

We first hypothesized that the *execute self-touch* condition would yield lower PSEs than the *execute posture* condition, replicating the classic phenomenon of sensory attenuation. Similar to Experiment 1, we further predicted that the *observe self-touch* condition would also yield lower PSEs than the *observe posture* condition, reflecting the attenuation during action observation due to simulation. Finally, by contrasting the *execute self-touch* and the *observe self-touch*, we would be able to compare the extent to which the action was simulated during action observation with respect to action execution. Any effects due to the mere presence of the actress would be captured by the comparison between the two control conditions: *execute posture* and *observe posture*. As in Experiment 1, we further checked for any sex effects given that the observed agent was always female.

### Experiment 3

#### Methods and Statistical analysis

Experiment 3 was identical to Experiment 2, with the only difference being that the *execute posture* and *observe posture* conditions were replaced by two new control conditions: *execute delayed self-touch* and *observe delayed self-touch* (**Figure 3**).

**Figure 3.**
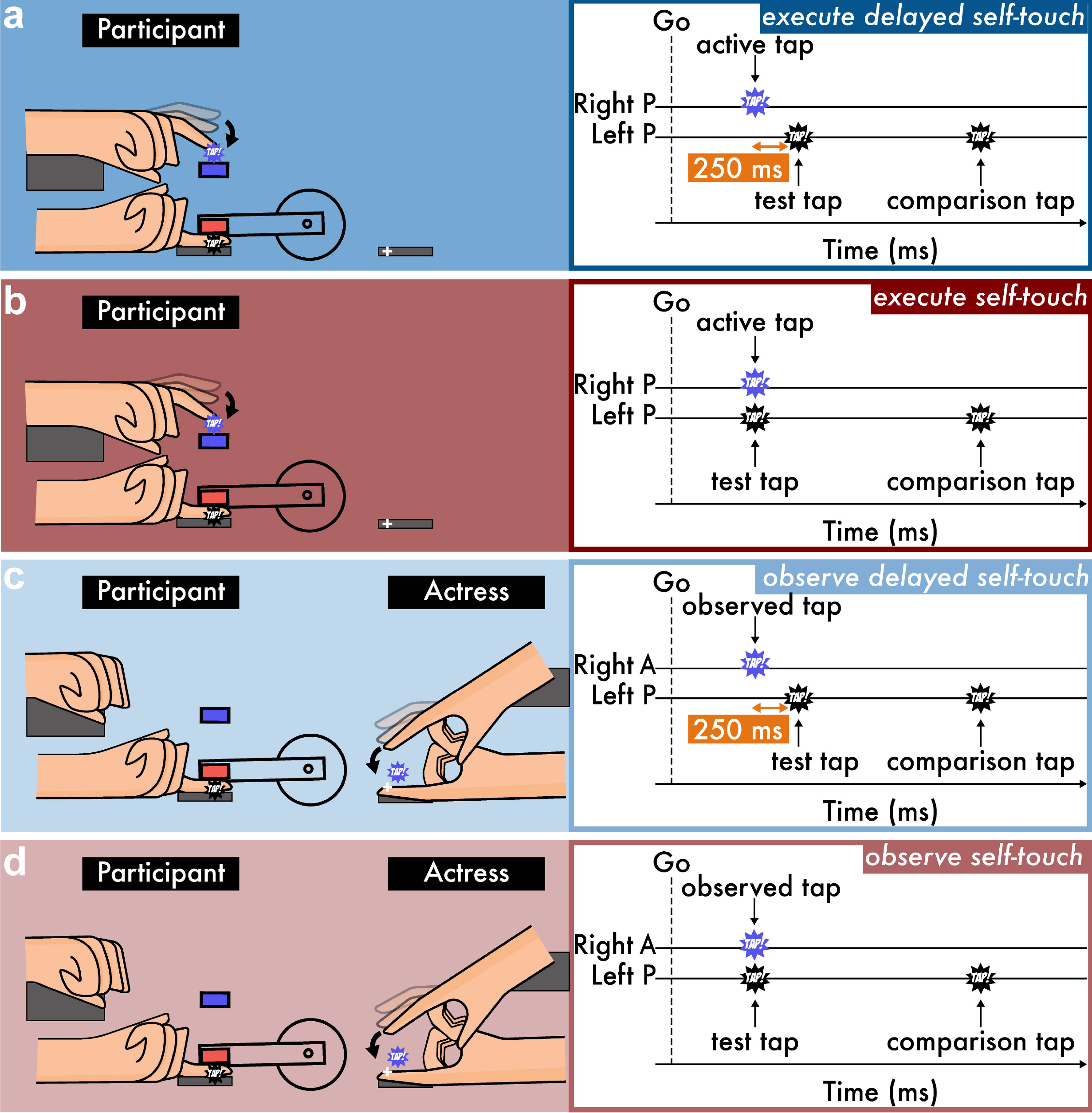
Experimental conditions in the force-discrimination task (Experiment 3). As in Experiment 2, participants received two taps on the pulp of their left index finger, and they verbally indicated which felt stronger: the first or the second tap. **(a)** In the *execute delayed self-touch* condition, the participants triggered the test tap by actively tapping with their right index finger a force sensor placed just above their left index finger. A 250-ms delay was introduced between the active tap and the test tap. **(b)** The *execute self-touch* condition was the same as the *execute delayed self-touch* condition but without the delay. **(c)** In the *observe delayed self-touch* condition, the actress triggered the test tap by actively tapping with her right index finger a hidden force sensor that was placed below her left index finger. A 250-ms delay was introduced between her active tap and the participants’ test taps. **(d)** The *observe self-touch* condition was the same as the *observe delayed self-touch* condition but without a delay. Fixation points are denoted by a cross (+). Right P and left P represent the right and left index fingers of the participant, respectively, while right A represents the right index finger of the actress.

The *execute delayed self-touch* condition (**Figure 3a**) was identical to the *execute self-touch* condition (**Figure 3b**), with the only difference being that we now introduced a 250-ms delay between the participants’ active tap and the applied test tap. Similarly, the *observe delayed self-touch* condition (**Figure 3c**) was identical to the *observe self-touch* condition (**Figure 3d**), with the difference that a 250-ms delay was introduced between the actress’s active tap and the participants’ applied test tap. Again, in both observation conditions, the participants passively observed the actress without moving, and the initial arm postures of the participant and the actress were similar across conditions.

In addition, we instructed the actress to apply a pressure of approximately 2 N on her left index finger during the observation conditions to match the 2 N test tap participants were simultaneously receiving on their index fingers. Forty-four trials (44 out of 6720, 0.65%) were rejected because either the intensity of the test tap (2 N) was not applied correctly or the responses were not recorded. All methods and statistical analyses were the same as those of Experiment 2.

#### Hypothesis

We expected that the *execute self-touch* condition would yield lower PSEs than the *execute delayed self-touch* condition, given previous results that, when a delay was introduced between the movement and its sensory consequences, the attenuation was greatly reduced because the stimulus no longer corresponded to the predicted consequence of the motor command (Blakemore et al. 1999; Bays et al. 2005; Kilteni et al. 2019). For this reason, the *executed delayed self-touch* condition served as a control condition. Similar to Experiments 1 and 2, we predicted that the *observe self-touch* would also yield lower PSEs than the *observe delayed self-touch* condition, reflecting the simulation-driven attenuation during action observation. The delayed self-touch condition served as a control condition in the observation session since it included the exact same movement of the actress but preceded the somatosensory stimulation by 250 ms. By contrasting the *execute self-touch* with the *observe self-touch*, we would be able to compare the extent to which the action was simulated during action observation with respect to action execution. Any effects due to the mere presence of the actress would be captured by the comparison between the two control conditions: *execute delayed self-touch* and *observe delayed self-touch*. As in Experiments 2 and 3, we further checked for any sex effects given that the observed agent was always female.

## Results

### Experiment 1

In Experiment 1, we investigated whether observing an actress reaching with her right index finger and pressing against her left index finger can influence the perceived magnitude of touches applied on the participants’ left index finger. We hypothesized that in this *observe self-touch* condition, the participants would reproduce weaker matched forces than in the *observe posture* condition, which does not include any action observation, and the *observe self-press* condition, which includes the observation of an action without somatosensory consequences for the left index finger. The results from Experiment 1 are summarized in **Figure 4**.

**Figure 4.**
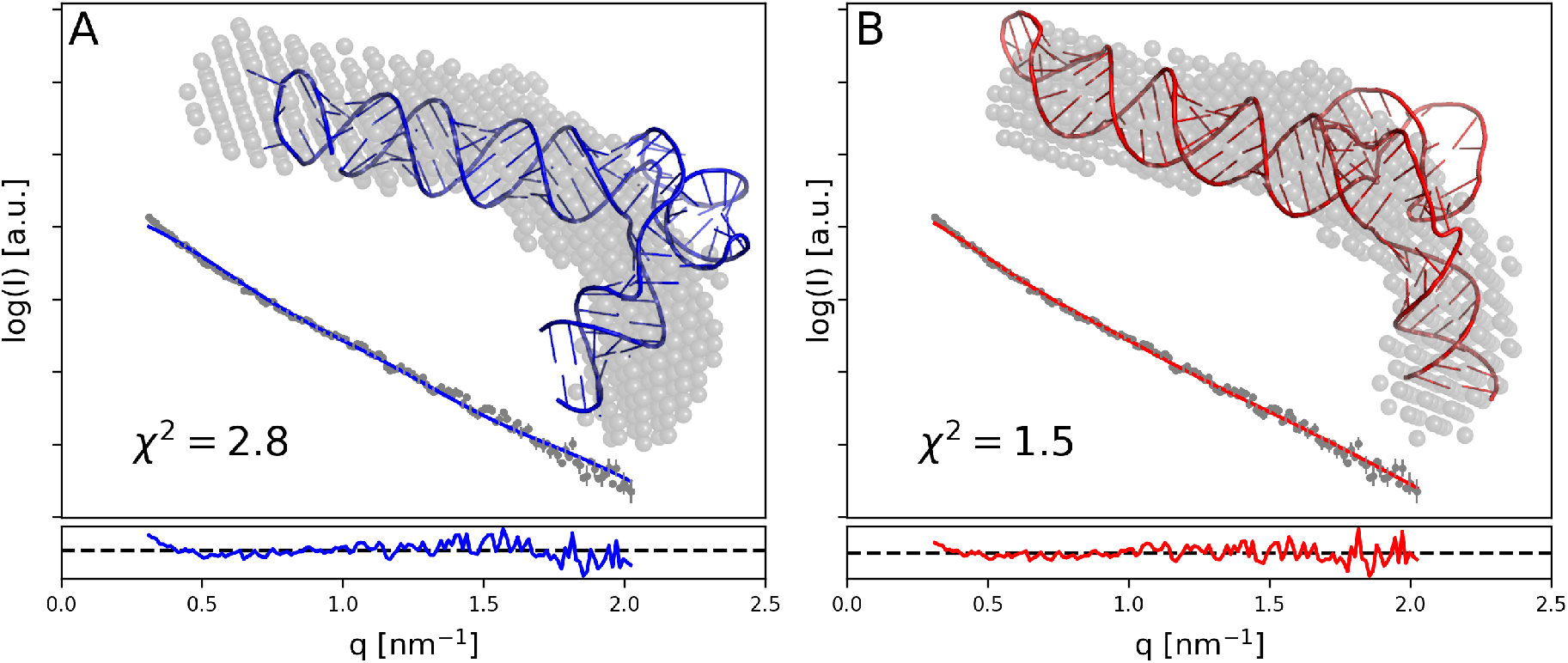
Results from the force-matching task of Experiment 1. **(a)** Forces (mean ± SE) generated by the participants (*matched* forces) as a function of the externally generated forces (*applied* forces). **(b)** Matched forces for each condition (mean ± SE) averaged across the applied force levels. **(c)** Forces (mean ± SE) generated by the actress (*observed* forces) as a function of the externally generated forces (*applied* forces). In **(a, c)**, the dotted lines indicate the theoretically perfect performance. Colored lines represent the fitted regression lines for each condition. For illustration purposes, the position of the markers has been scattered to avoid overlapping points. **(d, e, f)** Line plots illustrate the participants’ paired responses for each combination of conditions.

A repeated-measures ANOVA of the matched forces revealed a significant main effect of the applied force (*F*(5, 145) = 374.7, *p* < 0.001), a significant main effect of the condition (*F*(2, 58) = 3.456, *p* = 0.038) and no significant interaction between the applied force and the condition (*F*(10, 290) = 1.535, *p* = 0.126) (**Figure 4a**). Pairwise comparisons revealed significant differences between all pairs of applied force levels, confirming that the participants discriminated well the different force levels (all *p*-values < 0.001). With respect to the different conditions, the two control conditions, *observe posture* and *observe self-press*, did not significantly differ from each other (*t*(29) = −1.028, *p* = 0.312, *CI*^*95*^ = [−0.130, 0.043], *BF*_*01*_ = 3.175) (**Figure 4b,d-f**). Interestingly, the participants reproduced significantly weaker forces in the *observe self-touch* condition than the *observe self-press* condition (*t*(29) = −2.557, *p* = 0.016, *CI*^*95*^ = [−0.216, 0.024], *BF*_*01*_ = 0.330). At first, this difference could indicate that the participants predicted the somatosensory consequences of the *observe self-touch* condition; they attenuated their received touches and, therefore, reproduced weaker forces. However, the *observe self-touch* condition did not significantly differ from the *observe posture* condition (*t*(29) = −1.554, *p* = 0.131, *CI*^*95*^ = [−0.177, 0.024], *BF*_*01*_ = 1.749), which did not involve any action to observe. This absence of a significant difference speaks against a simulation process for the observed action within the motor system of the observer, but the Bayesian analysis provided only anecdotal evidence in favor of the null hypothesis (i.e., the absence of an effect was 1.749 times more likely than the existence of an effect given the data). Finally, the forces the actress pressed did not significantly differ between the *observe self-touch* and the *observe self-press* conditions (*t*(29) = −0.178, *p* = 0.860, *CI*^*95*^ = [−0.022, 0.018], *BF*_*01*_ = 5.068) and matched well with the applied forces (**Figure 4c**). No significant sex effects were observed (**Supporting Information**).

We did not find any conclusive evidence of somatosensory attenuation during action observation in Experiment 1. However, Experiment 1 did not include a condition where the participants actually performed self-generated touches in which we could verify the ‘normal’ attenuation of the participants driven by action execution. Including this condition would allow us to confirm that lack of attenuation driven by action observation occurred in combination with significant attenuation driven by action execution. Experiment 2 was designed to address this limitation. Since our design of the force-matching task did not allow such a condition because participants always used the slider – and not their finger – to reproduce the forces, Experiment 2 used the force-discrimination task, which quantified the attenuation of self-produced taps through discrimination rather than reproduction (Bays et al. 2005; Kilteni et al. 2019, 2020).

### Experiment 2

In Experiment 2, we investigated whether observing an actress reaching and tapping against her left index finger with her right index finger can influence the perceived magnitude of taps applied on the participants’ left index fingers using the force-discrimination task. We hypothesized that the participants would perceive the taps as having a weaker intensity (the PSEs would be lower) in the *execute self-touch* than in the *execute posture* condition, replicating the basic phenomenon of somatosensory attenuation. Similarly, the PSEs would be lower in the o*bserve self-touch* than in the *observe posture* condition, showing attenuation through simulation during action observation. With respect to JNDs, we did not hypothesize any specific differences between the two *self* conditions (*execute posture* and *execute self-touch*) based on our previous study (Kilteni et al. 2020). Similarly, we did not expect any differences between the two *observation* conditions (*observe posture* and *observe self-touch*). The results from Experiment 2 are summarized in **Figure 5**.

**Figure 5.**
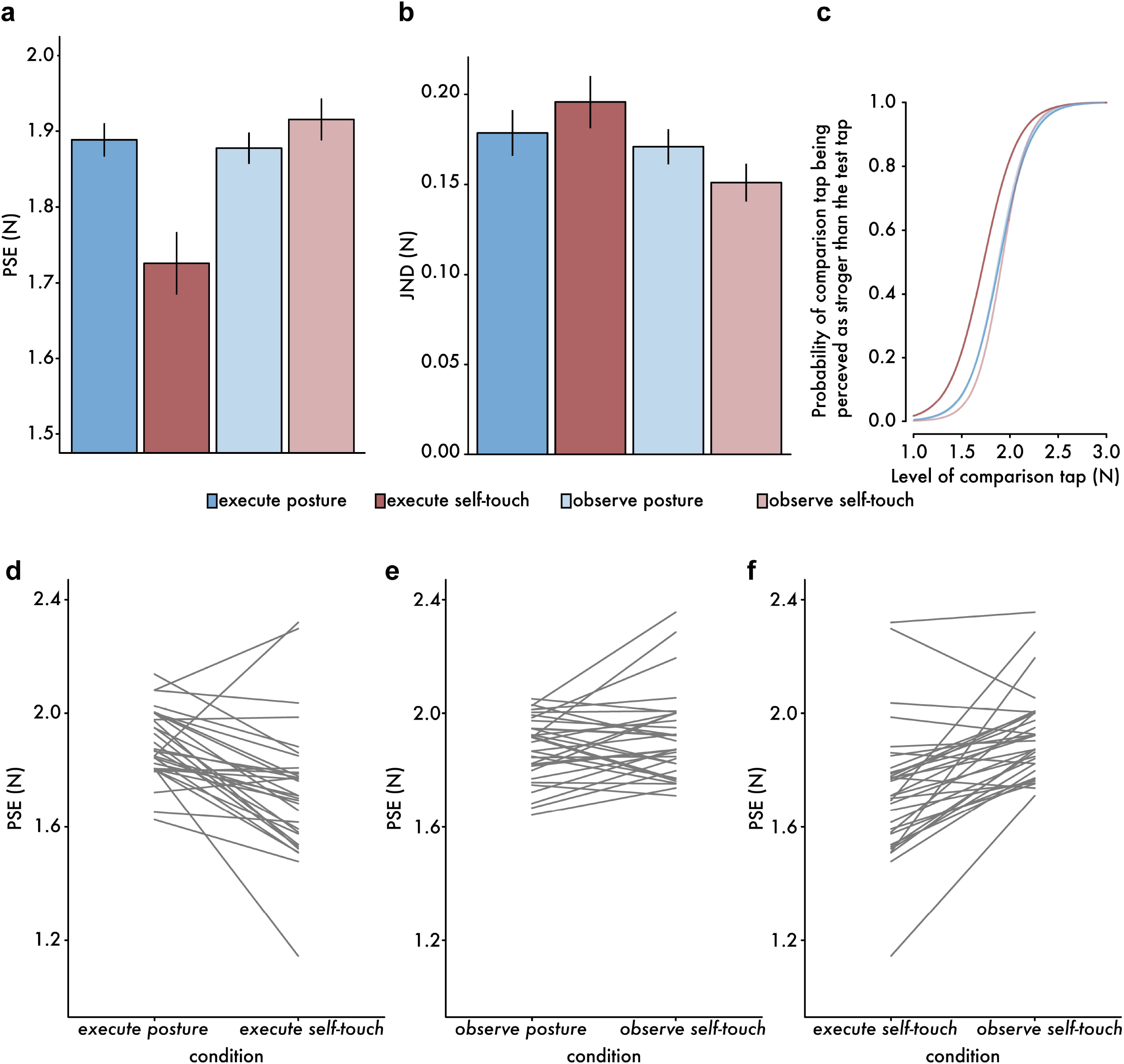
Results from the force-discrimination task of Experiment 2. **(a)** Bar graphs show the PSEs (mean ± SEM) for each condition. **(b)** Bar graphs show the JNDs (mean ± SEM) for each condition. **(c)** Group psychometric functions for each condition generated using the mean PSE and the mean JND across participants for the different levels of the comparison tap. The *execute posture* and *observe posture* conditions (blue curves) are almost completely overlapping since they had very similar PSE and JND parameters (**a, b**).**(d, e, f)** Line plots illustrate the participants’ paired responses for each combination of conditions.

As expected from previous self-touch studies, the PSEs were lower in the *execute self-touch* condition than in the *execute posture* condition (**Figure 5a, d**) (*t*(31) = −4.451, *p* < 0.001, *CI*^*95*^ = [−0.237, −0.088], *BF*_*01*_ = 0.004), replicating previous findings that a self-generated tap feels weaker than an externally generated identical tap (Bays et al. 2005; Kilteni et al. 2019, 2020). In contrast to the action observation motor simulation hypothesis, there was no significant difference between the *observe self-touch* condition and the *observe posture* condition (*t*(31) = 1.629, *p* = 0.113, *CI*^*95*^ = [−0.010, 0.085], *BF*_*01*_ = 1.614) (**Figure 5e**). In addition, lower PSEs were found in the *execute self-touch* condition than in the *observe self-touch* condition (*t*(31) = −5.453, *p* < 0.001, *CI*^*95*^ = [−0.260, −0.119], *BF*_*01*_ =0.0003) (**Figure 5f**), suggesting that an executed movement is more efficient in yielding somatosensory attenuation than an observed movement. Moreover, no differences were detected between the two control conditions (*execute posture* and *observe posture)*, suggesting that the mere view of the actress did not influence the perceived intensity of the received forces: *t*(31) = 0.647, *p* = 0.522, *CI*^*95*^ = [−0.024, 0.046], *BF*_*01*_ = 4.363). Finally, no significant sex effects were observed (**Supporting Information**).

With respect to JNDs (**Figure 5b**), there was no difference between the *execute self-touch* condition and the *execute posture* condition (*t*(31) = 1.098, *p* = 0.281, *CI*^*95*^ = [−0.015, 0.049], *BF*_*01*_ = 3.053); this suggests that the participants’ force-discrimination capacity on their left hands was not influenced by the movement of their right hand. In the observation conditions, the discrimination capacity was similar between the *observe self-touch* condition and the *observe posture* condition (*t*(31) = −1.790, *p* = 0.083, *CI*^*95*^ = [−0.043, 0.003], *BF*_*01*_ = 1.278). A significant difference was detected between the JNDs of the *execute self-touch* and *observe self-touch* conditions (*t*(31) = 3.502, *p* = 0.001, *CI*^*95*^ = [0.019, 0.071], *BF*_*01*_ = 0.042); this means that action observation improved the participants’ discrimination capacity compared to action execution. Finally, no JND differences were detected between the two control conditions (*execute posture* and *observe posture)*: *t*(31) = 0.541, *p* = 0.592, *CI*^*95*^ = [−0.021, 0.036], *BF*_*01*_ = 4.623. Sex effects were nonsignificant (**Supporting Information**).

**Figure 5c** shows the group psychometric curves for each condition. It is evident that attenuation is observed only in the *execute self-touch* condition.

For Experiment 1, the findings of Experiment 2 speak against the simulation theory of action observation. This finding is shown in the absence of a significant difference between the *observe self-touch* and the *observe posture* conditions and in the presence of a significant difference between the *executed self-touch* and the *observe self-touch* conditions. Nevertheless, we should admit that the Bayesian analysis remained inconclusive about the absence of a difference between the two conditions (*BF*_*01*_ = 1.614). Further, it is worth mentioning that we observed a significant improvement in the force-discrimination capacity of the participants when they observed the movement of the actress; this could indicate that observing a bimanual movement can increase the participants’ attention towards their own hands, which improves their sensitivity. Finally, the intensity of the taps the actress pressed (*mean* ± *sd*, 1.269 ± 0.266 N) was lower than the intensity of the taps the participants simultaneously experienced (2 N). This difference could have made the motor simulation more difficult. To rule out that our null findings were not due to any of the abovementioned reasons, we designed a new experiment that involved the execution and observation of the same movements in the control conditions as well, and the actress was then instructed to press exactly 2 N.

### Experiment 3

In Experiment 3, we hypothesized that the PSE would be lower in the *execute self-touch condition* than in the *execute delayed self-touch* condition, replicating previous findings on attenuation reduction due to the presence of a delay between the movement and its sensory consequence. Importantly, we further tested the prediction that PSEs should be lower in the o*bserve self-touch condition* than in the *observe delayed self-touch* condition based on the hypothesis that action observation involves motor simulation and sensory attenuation. With respect to JNDs, we expected no differences between the two *self* conditions (*execute self-touch* and *execute delayed self-touch*) or between the two *observation* conditions (*observe self-touch* and *observe delayed self-touch*). The results from Experiment 3 are summarized in **Figure 6**.

**Figure 6.**
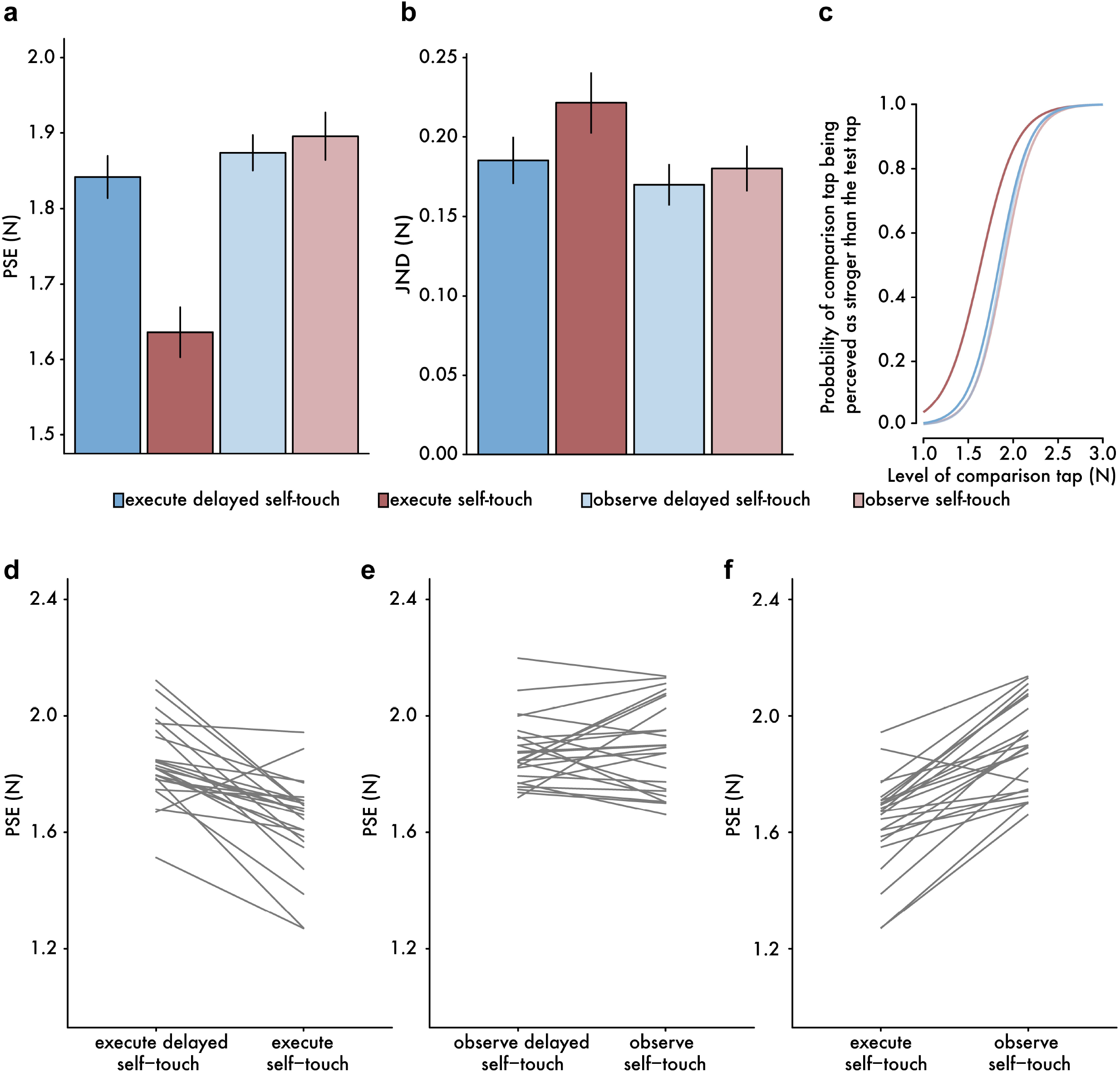
Results from the force-discrimination task of Experiment 3. **(a)** Bar graphs show the PSEs (mean ± SEM) for each condition. **(b)** Bar graphs show the JNDs (mean ± SEM) for each condition. **(c)** Group psychometric functions for each condition generated using the mean PSE and the mean JND across participants for the different levels of the comparison tap. **(d, e, f)** Line plots illustrate the participants’ paired responses for each combination of conditions.

As expected, the PSEs were lower in the *execute self-touch* condition than in the *execute delayed self-touch* condition (**Figure 6a, d**) (*t*(23) = −5.824, *p* < 0.001, *CI*^*95*^ = [−0.279, −0.133], *BF*_*01*_ < 0.001), replicating previous findings that a delayed self-generated tap feels stronger than an identical nondelayed self-generated tap (Bays et al. 2005; Kilteni et al. 2019). Nevertheless, there was no significant difference between the *observe self-touch* condition and the *observe delayed self-touch* condition, and the Bayesian analysis clearly supported the null hypothesis (*t*(23) = 0.792, *p* = 0.436, *CI*^*95*^ = [−0.035, 0.079], *BF*_*01*_ = 3.508) (**Figure 6e**). As in Experiment 2, lower PSEs were found in the *execute self-touch* condition than in the *observe self-touch* condition (*t*(23) = −8.511, *p* < 0.001, *CI*^*95*^ = [−0.323, −0.196], *BF*_*01*_ <0.001) (**Figure 6f**), suggesting again that an active movement is more efficient in yielding somatosensory attenuation than an observed movement. Moreover, no significant differences were detected between the two control conditions (*execute delayed self-touch* and *observe delayed self-touch)*: *t*(23) = −1.339, *p* = 0.194, *CI*^*95*^ = [−0.082, 0.017], *BF*_*01*_ = 2.112). Sex effects were nonsignificant (**Supporting Information**).

With respect to JNDs (**Figure 6b**), there was no difference between the *execute self-touch* condition and the *execute delayed self-touch* condition (*t*(23) = 1.729, *p* = 0.097, *CI*^*95*^ = [−0.007, 0.080], *BF*_*01*_ = 1.282). Similarly, in the observation conditions, no differences were detected between the *observe self-touch* condition and the *observe delayed self-touch* condition (*t*(23) = 0.581, *p* = 0.567, *CI*^*95*^ = [−0.026, 0.047], *BF*_*01*_ = 3.997). A statistical trend was detected between the JNDs of the *execute self-touch* and *observe self-touch* conditions (*n* = 24, *V* = 218, *p* = 0.053, *CI*^*95*^ = [−0.0004, 0.071], *BF*_*01*_ = 0.928), suggesting that action observation tended to improve the participants’ discrimination capacity. Nevertheless, the Bayesian analysis remained inconclusive, since it gave the same likelihood for the presence and the absence of an effect. Finally, no JND differences were detected between the two control conditions (*execute delayed self-touch* and *observe delayed self-touch)*: *t*(23) = 0.897, *p* = 0.379, *CI*^*95*^ = [−0.020, 0.051], *BF*_*01*_ = 3.244. Again, sex effects were nonsignificant (**Supporting Information**).

It is noteworthy that in Experiment 3, the intensity of the taps the actress pressed (*mean* ± *sd*, (2.064 ± 0.114 N) matched well the intensity of the taps the participants simultaneously experienced (2 N).

**Figure 6c** shows the group psychometric curves for each condition. As in Experiment 2, it is clear that attenuation is observed only in the *execute self-touch* condition.

## Discussion

In the present study, we investigated whether observing an agent reaching with her right hand to touch her left index finger would trigger the observer’s motor system to simulate the action and its somatosensory consequences and thus lead to the attenuation of a somatosensory stimulus simultaneously presented on the corresponding part of the observer’s left index finger. In three experiments, we did not find any reliable evidence for attenuation during action observation. Therefore, our data are not compatible with the direct-matching hypothesis, according to which the observer’s motor system automatically simulates the observed action. Below, we summarize our findings and discuss possible reasons for the absence of effects.

In Experiment 1, we used the classic force-matching task (Shergill et al. 2003; Bays and Wolpert 2008), and we compared the participants’ perceived intensity of pressure applied to their left index fingers during the observation of an actress reaching and touching her left index finger or the table with her right index finger or in a control condition when the actress held her right hand still (i.e., static posture). The perception of force during the observation of the reach and touch action did not significantly differ from that during the observation of the static posture. This finding speaks against the direct-matching hypothesis, since a simulation is expected in the former case but not in the latter case. Nevertheless, the Bayesian analysis did not conclusively support the absence of any effects. Furthermore, we observed that participants felt the touch significantly weaker when observing the reaching movement to touch the left index finger than when observing the reaching movement to touch the table. This difference could provide some evidence for the direct-matching hypothesis, since different simulations would be expected for different movements. However, somatosensory perception during the observation of the reaching movement to touch the table did not significantly differ from that during posture observation, with the Bayesian analysis now supporting the absence of effects. When taken together, these results do not provide conclusive support in favor of or against the direct-matching hypothesis. Therefore, we designed Experiment 2 to re-address our research question with new conditions and a different task.

In Experiment 2, we used a force-discrimination task, and we compared the participants’ perceived intensity of a somatosensory stimulus applied to their left index fingers during the observation of an actress reaching and touching her left index finger with her right index finger or staying still (i.e., static posture). We further included two conditions that involved actual execution of the same reaching movement or staying still (i.e., static posture). By this approach, we were able to directly compare attenuation during action execution with attenuation during action observation. We found significant attenuation during action execution compared to posture execution, in line with previous research (Bays et al. 2005; Kilteni et al. 2019, 2020). Importantly, we found no significant attenuation during action observation compared to posture observation. This null finding speaks against the direct-matching hypothesis, since a simulation is expected in the former but not in the latter condition. However, the Bayesian analysis did not conclusively support the absence of any effects. In addition, participants perceived their self-generated touches to be significantly weaker when executing the action than when observing the action. This further speaks against the direct-matching hypothesis, since a similar recruitment of the observer’s motor system is theoretically expected during both action execution and action observation. Furthermore, we observed that participants had a significantly better discrimination capacity when observing the action of the actress than when executing the action themselves, and this improvement could have confounded our findings with different attention requirements in the experimental conditions. Therefore, we designed Experiment 3 to further examine our research question with very well-matched conditions.

In Experiment 3, we again used the force-discrimination task, and we compared the participants’ perceived intensity of a somatosensory stimulus applied to their left index fingers during the execution or observation of a reaching movement to touch the left index finger. In the control conditions, we included a delay between the executed/observed movement and the resulting touch. By doing so, we were able to directly compare action observation and action execution conditions that all involved movement, which is an advantage with respect to Experiment 2, where passive conditions were used as controls. We found significant attenuation during action execution compared to delayed action execution, in line with previous research (Blakemore et al. 1999; Bays et al. 2005; Kilteni et al. 2019). However, the participants’ performance during action observation and delayed action observation did not significantly differ. The Bayesian analysis further supported the absence of any effects. Once again, this finding speaks against the direct-matching hypothesis. Furthermore, and in line with the findings from Experiment 2, participants perceived their touches to be significantly weaker when executing the nondelayed action than when observing the nondelayed action. This again speaks against the direct-matching hypothesis, since the engagement of the observer’s motor system is expected during both action execution and action observation. Taken together, the results of Experiment 3 constitute conclusive evidence against the direct-matching hypothesis. Thus, at least during the action observation conditions used in the current study, the participants did not automatically simulate the observed movement to the extent that this produced the attenuation of somatosensory stimuli applied to their body.

A few earlier studies on sensory attenuation during action observation exist, but these have yielded mixed results. In his study, Sato (2008) observed that participants perceived the loudness of a sound as weaker both when they generated it by pushing a button and when they observed the experimenter performing the same action. Indeed, this specific result would be expected according to the simulation theory. However, no such effects were observed in the subsequent study of Weiss et al. (2011), who observed auditory attenuation only when the subjects produced the sound themselves and not when observing the experimenter producing it. Rather than attenuation, Thomas et al. (2013) observed perceptual enhancement during action observation and proposed that the observer simulates the sensations of the observed agent and not the motor program. They suggested that this enhancement could relate to findings that watching somebody being touched (Keysers et al. 2004) or manipulating an object (Avikainen et al. 2002) activates the observer’s somatosensory cortex. Our study did not reveal sensory attenuation or enhancement during action observation; in contrast, only self-generated tactile stimuli were attenuated, while those delivered during action observation were processed as if they were fully externally generated, which, of course, they actually were.

Why did we not observe any attenuation effects during action observation, and why are earlier results not consistent? One explanation is that the function of mirror neurons is not action understanding through simulation, as has been argued (Rizzolatti and Craighero 2004; Rizzolatti and Sinigaglia 2010; Giese and Rizzolatti 2015). Criticism has been raised about the simulation theory (Saxe 2005) and on whether there is evidence showing that monkeys understand the observed action (Oztop et al. 2006, 2013; Hickok 2009). Moreover, the multiple interpretations given to the notions of motor resonance and action/goal understanding have brought further uncertainty (Uithol et al. 2011; Cook et al. 2014): for example, it is not clear whether it is the observed action, the goal of the observed action, or the action that is anticipated in response to the observed action that should be understood by mirror neurons (Uithol et al. 2011). A strict interpretation of this action understanding functionality would dictate that mirror neurons are selective in the actions they encode and that they respond to the same type of movement during both execution and observation (Dinstein et al. 2008). However, this is not always the case: not all mirror neurons are selective to only one type of action, and a large proportion of mirror neurons discharge for observed and executed actions that have different goals (Csibra 2005; Hickok 2009). In light of these observations, alternative views proposed that mirror neurons anticipate the subsequent actions of the observed agent (Csibra 2005) or reflect task-dependent sensory–motor associations (Hickok 2009; Heyes 2010a, b). Therefore, if the activity of the putative human mirror system is not to automatically simulate what we see, the absence of sensory attenuation in our study should not be surprising.

It can also be argued that the direct hypothesis, and thus the automatic simulation, apply only for certain types of movements, that is, those that involve meaningful hand-object interactions. In our experiments, participants observed a person reaching to tap her left index finger with her right index finger. This is a simple task that is present in the observers’ motor repertoire (Giese and Rizzolatti 2015), and that does not require complex learned kinematics. However, it is an intransitive action, i.e., an action that does not involve any object grasping, object manipulation, or holding of an external object. In primates, both transitive (object-directed)(Rizzolatti and Craighero 2004) and intransitive movements (Kraskov et al. 2009) are effective in eliciting the activation of the mirror neurons (see also (Cook et al. 2014)). Similarly, in humans, experimental evidence suggests that both transitive and intransitive movements can elicit activity in the putative human mirror neuron system (Fadiga et al. 1995; Jonas et al. 2007; Lui et al. 2008). Therefore, theoretically speaking, one should expect that our participants would simulate the observed action and attenuate the external touches, even if the observed movement is relatively simple and not involving an external object.

A defender of the direct-matching hypothesis could further postulate that the simulation is not sufficiently accurate to allow the full engagement of internal models that compute the sensorimotor predictions; in other words, there is an automatic simulation process during action observation but at a more abstract high level, or there is an automatic simulation that engages the internal models to a lesser extent than action execution. Although the mirror neurons have been explicitly related to predictive processing (Kilner et al. 2007a, b; Urgen and Miller 2015) and internal forward models (Miall 2003; Wolpert et al. 2003; Oztop et al. 2006, 2013; Imamizu 2010), the defender could also argue that during action observation, the observer performs inverse modeling, i.e., the mapping of the visual representation of the action to the motor program, but not forward modeling, i.e., the mapping of the motor program to its sensory consequences. It is interesting to note that a recent meta-analysis comparing networks of action execution and action observation revealed no activation of the cerebellum (Hardwick et al. 2018). The cerebellum has been proposed to host internal forward models and support motor prediction (Miall and Wolpert 1996; Wolpert et al. 1998; Shadmehr and Krakauer 2008). Previous studies on somatosensory attenuation have revealed cerebellar activity during self-generated touches compared to externally generated touches (Blakemore et al. 1998; Shergill et al. 2013; Kilteni and Ehrsson 2020a), while a recent study revealed that the functional connectivity between the cerebellum and the primary and secondary somatosensory cortexes reflects somatosensory attenuation at the behavioral level (Kilteni and Ehrsson 2020a). In contrast to action observation, motor imagery, i.e., imagining the execution of an action without performing it, recruits the cerebellum (Lotze et al. 1999; Grezes and Decety 2001; Hardwick et al. 2018) and produces sensory attenuation (Kilteni et al. 2018) similar to motor execution. Therefore, we could speculate that the absence of cerebellar activation during action observation might indicate that the observer does not, or cannot, predict the consequences of the observed action (forward modeling) but can derive the motor command from the visual representation of the action (inverse modeling). In this case, no attenuation should be expected.

Finally, one can argue that the action observation effects are very small and difficult to detect with our behavioral methods. In monkeys, the activity of mirror neurons in the primary motor cortex is substantially weaker during observation than execution (Dushanova and Donoghue 2010; Vigneswaran et al. 2013; Kraskov et al. 2014; Jerjian et al. 2020). Moreover, there exist mirror neurons that discharge during action execution but are actually suppressed during action observation (Kraskov et al. 2009; Vigneswaran et al. 2013; Jerjian et al. 2020). In humans, electrophysiological responses to median nerve stimulation are suppressed to a greater extent during action execution than action observation (Hari et al. 1998), the spectral power is differentially modulated for action observation and action execution (Cochin et al. 1999; Silas et al. 2010) (see also (Waldert et al. 2015)) and the corticospinal excitability effects observed during action observation (Fadiga et al. 1995; Gueugneau et al. 2015) might be weaker or less related to action execution (Bunday et al. 2016; Hannah et al. 2018) than theorized by the direct-matching hypothesis. In the present study, we quantified somatosensory attenuation, a robust phenomenon manifested during voluntary action (Bays et al. 2005; Kilteni et al. 2020) that is theorized to result from the predictions of the internal forward models (Wolpert and Ghahramani 2000; Blakemore et al. 2000; Wolpert and Flanagan 2001; Bays and Wolpert 2008) and not from generalized gating processes (Kilteni and Ehrsson 2020b). We have used this specific experimental approach in a previous study that investigated whether motor imagery includes simulation processes similar to those during action execution (Kilteni et al. 2018). In that study, participants were explicitly asked to imagine pressing one index finger against the other, and we observed that this motor imagery produced an attenuation of the somatosensory stimuli applied on the same finger they imagined that they were touching (Kilteni et al 2018). Thus, if action observation engages a similar internal simulation process as action execution and mental motor imagery, the present paradigm should be able to detect it as a force attenuation effect. In addition, in all three experiments (Experiments 1-3), we included control conditions that always involved the actress to account for additional confounds that otherwise could hinder the attenuation during action observation. For example, watching the actress’s left index finger might increase the attention to one’s own left index finger and bias perception; therefore, this factor was controlled for by including an action observation control condition with similar visual input. Moreover, we further controlled that participants observed similar actions in the experimental and control conditions (Experiment 3) and that the forces they observed were congruent with the forces they received (Experiment 1, Experiment 3). Furthermore, a post hoc power sensitivity analysis revealed that our experiments were sensitive enough to reliably detect any medium effect size with 80% power (paired-samples t-tests, *alpha* = 0.05, Cohen’s *d* =[0.511, 0.597]). Unless the effects were smaller than our minimum detectable effect sizes, we consider that we would have observed some somatosensory attenuation effects if there was a consistent simulation of the observed action within the observer’s motor system.

In the present study, we tested one of the key suggested functions of the mirror neuron system; that we automatically simulate observed actions to even the extent that we predict their sensory consequences; a fundamental component of predictive sensorimotor control. When considering the results from our three experiments as a whole, we found systematic evidence for somatosensory attenuation during action execution but no signs of attenuation during action observation. Distinguishing (and attenuating) self-generated from externally generated somatosensory sensations is crucial for our survival, because we need to discriminate the touches coming from our self from those coming from other individuals and animals, e.g., a predator. If we predicted all actions we observed and attenuated all their somatosensory consequences, we could potentially mistake externally generated sensations for self-generated ones and expose ourselves to risks that could threaten our health and safety. Next, the attenuation of sensory signals during action observation to the same degree as during action execution could constitute a limitation of the motor system rather than an advantage. Our results emphasize the tight link between somatosensory attenuation and self-generated actions and thus suggest that the basic distinction between self and others is maintained in the central motor system during the observation and execution of an action.

## Acknowledgments

K.K. was supported by the Swedish Research Council (VR Starting Grant, registration number 2019-01909). This work was supported by the Swedish Research Council (Distinguished Professor Grant, registration number 2017-03135, to H.H.E.).

## Data Accessibility

Data from all experiments are accessible upon request.

## Author Contributions

K.K. and H.H.E. conceived and designed the experiments. K.K. and P.E. collected the data of Experiment 1 and Experiment 2; specifically, K.K. was the actress and P.E. collected the force responses. I.B. and L.M. collected the data from Experiment 3; specifically, L.M. was the actress and I.B. collected the force responses. K.K. conducted the statistical analyses. K.K. and H.H.E. wrote the manuscript. P.E., I.B. and L.M. read and approved the final version.

