## Supporting information for "No evidence for somatosensory attenuation during action observation of self-touch"

**Sex effects**

We did not hypothesize any sex effects in any of the three experiments, as we reasoned that any automatic simulation of the observed action by the putative human mirror neuron system should not depend on the sex of the observed agent. Nevertheless, we statistically tested for any influences of the participants' sex since the actress was always female. To do so, we created a categorical factor, 'sex', representing the participants' sex, and we performed the main analysis of each experiment by including this additional factor.

Experiment 1

In Experiment 1, we performed a mixed ANOVA with the applied force and the condition as the within-subjects factors and sex as the between-subjects factor. The mixed ANOVA revealed a nonsignificant main effect of sex ($F(1, 28) = 0.035, p = 0.853$) and nonsignificant interactions between sex and applied force ($F(5, 140) = 0.462, p = 0.804$), between sex and condition ($F(2, 56) = 1.361, p = 0.265$) and among sex, applied force and condition ($F(10, 280) = 1.146, p = 0.328$).

Experiment 2

In Experiment 2, we performed a mixed ANOVA with condition as the within-subjects factor (*execute posture*, *execute self-touch*, *observe posture*, *observe self-touch*) and sex as the between-subjects factor. With respect to the PSEs, there was no significant main effect of sex ($F(1, 30) = 0.187, p = 0.668$) and no significant interaction between sex and condition ($F(3, 90) = 1.356, p = 0.262$). With respect to the JNDs, there was no significant main effect of sex ($F(1, 30) = 0.645, p = 0.428$) and no significant interaction between sex and condition ($F(3, 90) = 0.673, p = 0.571$).

Experiment 3

In Experiment 3, we performed a mixed ANOVA with condition as the within-subjects factor (*execute self-touch*, *execute delayed self-touch*, *observe self-touch*, *observe delayed self-touch*) and sex as the between-subjects factor. With respect to the PSEs, there was no significant main effect of sex ($F(1, 22) = 3.89, p = 0.061$) and no significant interaction between sex and condition ($F(3, 66) = 0.896, p = 0.448$). With respect to the JNDs, there was no significant main effect of sex ($F(1, 22) = 0.494, p = 0.490$) and no significant interaction between sex and condition ($F(3, 66) = 1.064, p = 0.371$).
